# Recombinational repair of nuclease-generated mitotic double-strand breaks with different end structures in yeast

**DOI:** 10.1101/2020.07.29.226639

**Authors:** Dionna Gamble, Samantha Shaltz, Sue Jinks-Robertson

**Author notes:** **Corresponding author:** Sue Jinks-Robertson, Department of Molecular Genetics and Microbiology, 213 Research Dr., Box 3020 DUMC, Duke University Medical Center, Durham, NC 27710.

## Abstract

Mitotic recombination is the predominant mechanism for repairing double-strand breaks in *Saccharomyces cerevisiae*. Current recombination models are largely based on studies utilizing the enzyme I-*Sce*I or HO to create a site-specific break, each of which generates broken ends with 3’ overhangs. In this study sequence-diverged ectopic substrates were used to assess whether the frequent Pol δ-mediated removal of a mismatch 8 nucleotides from a 3’ end affects recombination outcomes and whether the presence of a 3’ *versus* 5’ overhang at the break site alters outcomes. Recombination outcomes monitored were the distributions of recombination products into crossovers *versus* noncrossovers, and the position/length of transferred sequence (heteroduplex DNA) in noncrossover products. A terminal mismatch that was 22 nucleotides from the 3’ end was rarely removed and the greater distance from the end did not affect recombination outcomes. To determine whether the recombinational repair of breaks with 3’ *versus* 5’ overhangs differs, we compared the well-studied 3’ overhang created by I-*Sce*I to a 5’ overhang created by a ZFN (Zinc Finger Nuclease). Initiation with the ZFN yielded more recombinants, consistent with more efficient cleavage and potentially faster repair rate relative to I-*Sce*I. While there were proportionally more COs among ZFN-than I-*Sce*I-initiated events, NCOs in the two systems were indistinguishable in terms of the extent of strand transfer. These data demonstrate that the method of DSB induction and the resulting differences in end polarity have little effect on mitotic recombination outcomes despite potential differences in repair rate.

## INTRODUCTION

Double-strand breaks (DSBs) are one of the most toxic types of DNA damage that occur within the cell. These lesions can be caused by reactive oxygen species from DNA metabolism, collapsed or stalled replication forks, ionizing radiation, or other DNA damaging agents. If left unrepaired they can trigger loss of genetic information, genome instability and cell death. There are two major pathways for repairing DSBs in eukaryotes: nonhomologous end joining (NHEJ) and homologous recombination (HR). NHEJ directly re-ligates broken ends and this can be accompanied by sequence alterations at the joints or generate genome rearrangements if the ends of different breaks are joined. By contrast, HR is a relatively error-free process that uses a homologous sequence as a template to repair the break. Defects in HR have been linked to a wide range of human diseases that include cancer, neurodegenerative disorders and developmental disorders (Thompson and Schild 2002; Jackson and Bartek 2010; Hou *et al*. 2017).

HR mechanisms and pathways have been largely defined using the budding yeast *Saccharomyces cerevisiae* as a model system (Pâques and Haber 1999; Symington *et al*. 2014). All begin with 5’-end resection to generate long, 3’ tails that are assembled into nucleoprotein filaments that search for a homologous donor to be used as a repair template (Figure 1) (Cejka 2015; Symington 2016). End resection occurs in two steps. First, the Mre11-Rad50-Xrs2 (MRX) complex together with Sae2 initiates short-range resection that removes up to 200 nt. This is followed by long-range resection that can extend for thousands of nucleotides and is carried out by either the Exo1 exonuclease or the combined activities of the Sgs1 helicase (as part of the STR complex comprised of Sgs1, Top3 and Rmi1) and Dna2 endonuclease. Once a repair template is found, the 3’ tail pairs with the complementary strand of the homologous donor duplex, displacing the other strand to form a D (displacement) loop (San Filippo *et al*. 2008; Symington *et al*. 2014). The initial pairing of sequences from different duplexes creates a region of heteroduplex DNA (hetDNA). The invading 3’ end is then extended by a DNA polymerase, most likely Pol δ (Li *et al*. 2009; Guo *et al*. 2017; Donnianni *et al*. 2019). In the canonical double-strand break repair (DSBR) pathway, DNA synthesis expands the D-loop until sequence complementary to the 3’ end on the other side of the break is revealed. Pairing of the D-loop with the other end creates a second patch of hetDNA and double Holliday junctions (dHJs) are formed that can be dissolved or cleaved. Cleavage of the dHJs produces either noncrossover (NCO) or crossover (CO) products that have the linkages of sequences that flank the DSB maintained or switched, respectively. If the D-loop is dismantled before second-end engagement, the extended 3’ end anneals to the other side of the break, which creates a new patch of hetDNA, in a pathway termed synthesis-dependent strand annealing (SDSA). Subsequent gap filling and ligation yields an NCO product that contains a single tract of hetDNA (blue box in Figure 1) that can be either upstream (promoter-proximal) or downstream (promoter-distal) of the initiating DSB, depending on which end invades the donor duplex. By contrast, dissolution generates an NCO product with a tract of hetDNA on both sides of the initiating break.

**Figure 1.**
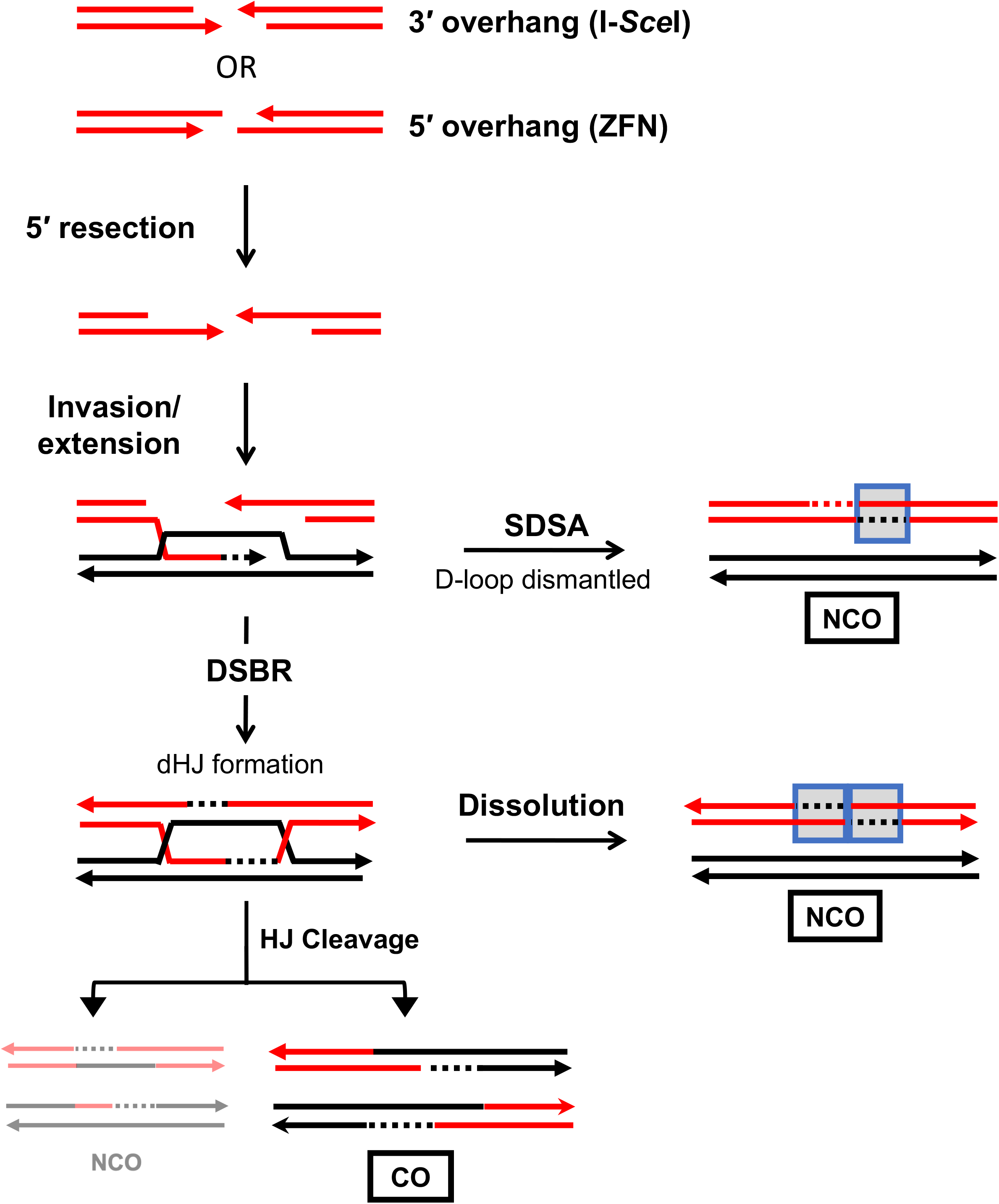
Repair of DSBs by the DSBR and SDSA pathways. Red and black lines represent the single strands of recipient and donor alleles, respectively, and arrowheads correspond to 3’ ends. Blue boxes indicate hetDNA and dotted lines are newly synthesized DNA that is the same color as the complementary template. The 5’ ends of a break are resected to generate 3’ tails that invade the donor. Upon strand invasion, a D-loop is formed and DNA synthesis extends the 3’ end. In the DSBR model, synthesis leads to second-end capture in which the displaced strand of the D-loop anneals to the other 3’ end of the break and dHJs are formed. dHJ dissolution produces a NCO product with hetDNA on each side of the initiating break (the donor is unchanged) while cleavage results in hetDNA on opposing sides of the break position in each CO or NCO product. During SDSA, the D-loop is dismantled and the newly synthesized 3’ end anneals to the other end of the break. The resulting NCO has hetDNA on only one side of the initiating break. In this system, NCOs generated by dHJ cleavage are very rare.

Molecular studies of mitotic recombination following the creation of a site-specific DSB have relied on the enzyme HO or I-*Sce*I, which makes DSBs with 4-nt, 3’ overhangs (Haber 1995, 2000; Sugawara and Haber 2006). DSBs that arise spontaneously, however, have end structures that potentially vary in overhang polarity, sequence and length. For example, when a replication fork encounters a nick, a DSB with a blunt end or overhang (3’ or 5’) can be generated depending on how the collision occurs (Morimatsu and Kowalczykowski 2014). In addition to “clean” end structures with terminal 5’-phosphate and 3’-hydroxyl groups, the ends of damage-induced spontaneous breaks may contain attached proteins or chemical groups, which must be removed before repair can be initiated and/or completed (Wyman and Kanaar 2006).

Several prior studies suggest that end structure may affect the subsequent repair process. An early example came from expression of restriction enzymes in yeast where DSBs created by *Eco*RI, which generates 5’ overhangs, were efficiently repaired and well tolerated (Lewis *et al*. 1999) while those generated by enzymes that created blunt ends were not (Westmoreland *et al*. 2010). A more recent study examined the effect of DSB overhang polarity and sequence on NHEJ (Liang *et al*. 2016). The HO endonuclease was used to generate DSBs with 3’ overhangs while those with 5’ overhangs were created using a Zinc Finger Nuclease (ZFN). The type of DSB affected protein recruitment, repair efficiency, and the choice between direct re-ligation or limited end modification before rejoining. Although NHEJ proteins were recruited more quickly to DSBs with 5’ overhangs and repair was more efficient, repair of DSBs with 3’ overhangs occurred with higher fidelity. The latter difference was attributed to 5’-end processing and 3’-end protection by the MRX complex. In addition to *in vivo* studies, the relationship between end structure and processing has been examined using purified proteins. The *Escherichia coli* RecJ protein, which is a 5’>3’ exonuclease on single-strand DNA, was more active on duplex DNA with 5’ overhangs than on 3’ overhangs or blunt ends (Morimatsu and Kowalczykowski 2014). This polarity was the reverse of that observed with yeast Exo1, which also is a 5’>3’ exonuclease. Exo1 had greater activity on duplex DNA with 3’ tails than with 5’ tails (Cannavo *et al*. 2013).

In the current study we investigated whether 3’ end removal by Pol δ prior to end extension or whether the enzyme used to create a site-specific DSB affected subsequent repair by the HR pathway. A common set of ectopic substrates was used and a recombination-initiating DSB was created using I-*Sce*I or a ZFN, which generate breaks with 4-nt 3’ overhangs or 4-nt 5’ overhangs, respectively. We examined the repair frequency, the CO-NCO distribution among products and lengths of hetDNA in NCO products. Although ZFN-induced breaks produced more recombinants and a slight increase in the proportion of COs, there were no significant differences in the NCO hetDNA profiles. Physical analysis highlighted different cleavage efficiencies between initiating enzymes and potential differences in repair rates.

## MATERIALS AND METHODS

### Media and Growth Conditions

Strains were stored at −80° and all growth was at 30°. YEPD (1% Bacto-yeast extract, 2% Peptone, 2% dextrose; 1.5% agar for plates) supplemented with 500 μg/ml adenine hemisulfate was used for nonselective growth. Drug-resistance markers were selected by adding the relevant drug to YEPD. For galactose-induction experiments, cells were grown in YEP supplemented with 2% raffinose (YEPR) instead of glucose. Synthetic complete (SC) medium contained 2% dextrose and was supplemented with all amino acids, uracil and adenine. For selection of prototrophs, the relevant amino acid or base was omitted (e.g., SC-Lys). Constitutive induction experiments required synthetic complete medium containing 2% galactose instead of dextrose to induce expression of the given nuclease. Ura^-^ segregants were selected on 5FOA plates (SC-Ura supplemented with 0.5 g uracil and 1 g 5-fluoroorotic acid per liter).

### Yeast strain construction

Lists of strains and relevant plasmids are provided in Tables S1 and S2, respectively. All strains were derived from W303 backgrounds by transformation or mating. Each haploid experimental strain contained a nuclease-cleavable recipient allele at *LYS2* (*hisG-lys2::NUC*), a nuclease-noncleavable donor allele at *CAN1* (*can1::lys2Δ3’::NUCnc-URA3-hisG*), and a galactose-regulated nuclease. Two independent isolates of each haploid were used in experiments. The insertion of a bacterial *hisG* gene upstream and a *URA3-hisG* cassette downstream of the recipient and donor alleles, respectively, conferred an unstable Ura^+^ phenotype for CO events.

The break-proximal SNPs in the original I-*Sce*I system (SJR3848) were 8 nt from the enzyme-created 3’ ends (SNP8) and frequently removed by Pol δ proofreading activity (Guo *et al*. 2017; Hum and Jinks-Robertson 2019). The *delitto perfetto* method (Storici and Resnick 2006) was used to move the break-proximal SNPs to 22 nt from the 3’ ends (SNP22). The CORE-UK cassette (*kanMX* and *URA3Kl* as selectable and counterselectable markers, respectively) was introduced into a donor-only strain (SJR4016) near the *BgI*II site in the *lys2Δ*3’ allele by selecting Ura^+^, G418-resistant colonies. The resulting strain (SJR4029) was then transformed with an amplified, 180-bp gBlock fragment (Integrated DNA Technologies) that had a non-cleavable I-*Sce*I site (I-SceInc) and contained SNP22 instead of SNP8 on each side of the break site. Ura^-^ transformants were selected and screened for *kanMX* loss and the desired sequence changes, yielding strain SJR4686. A *URA3-hisG* cassette was amplified from pNKY51 (Alani *et al*. 1987) and inserted downstream of the donor allele in SJR4686, creating SJR4727. SJR4727 was then crossed with SJR3848 to derive SJR4748, an *mlh1Δ* strain containing the *lys2::I-SceI* recipient allele, the donor *lys2Δ3’::I-SceIncSNP22-URA3-hisG* allele, and galactose-inducible I-*Sce*I. Finally, SJR4748 was crossed with SJR4258 (Hum and Jinks-Robertson 2019) to introduce a recipient allele with an upstream *hisG* marker (SJR4815).

Integrating plasmids containing the components of the ZFNs (pSR1109 and pSR1110) were adapted from replicating plasmids ryA-ZFN and ryB-ZFN (Beumer *et al*. 2006). *GAL1*-regulated ryA-ZFN and ryB-ZFN fragments were inserted into *Xho*I/*Sac*I-digested pRS303 and pRS305, which contain *LEU2* and *HIS3* as selectable markers, respectively (Sikorski and Hieter 1989). Following linearization, a single copy of each plasmid was inserted at *LEU2* or *HIS3*. The break-proximal SNPs in the initial ZFN chromosomal system were 7 nt from each 3’ end (SJR4582). To relocate the SNPs to 22 nt from the 3’ ends (SNP22), we utilized the same approach described above. SJR4029 was transformed with a 179-bp gBlock fragment containing a non-cleavable ZFN site (ZFNnc) and SNP22 on each side of the break, creating SJR4813. The *URA3-hisG* cassette was then amplified and inserted downstream of the donor in SJR4813 to create SJR4814. Finally, SJR4814 was crossed with SJR4582 to derive SJR5022, an *mlhl1Δ* strain containing the recipient *hisG-lys2::ZFN* allele, the donor *lys2Δ3’::ZFNncSNP22-URA3-hisG* allele, and galactose-inducible ZFN.

### DSB Induction

Strains were streaked to single colonies on YEPD and grown for two days. To measure repair (Lys^+^) frequencies, single colonies were inoculated into YEPR cultures and grown to an OD of 0.7-1.0 before plating on YPD and on selective plates supplemented with 2% galactose and lacking lysine. To limit enzyme cutting for subsequent genetic/molecular analyses (pulse inductions) single colonies were inoculated into individual 5 ml YEPR cultures and grown to an OD of 0.7-1.0. Galactose was added to 0.1% to induce expression of the relevant nuclease and incubation was continued for 45 min, which largely limited DSB formation to only one sister chromatid (Guo *et al*. 2017). Following induction, cells were washed and appropriate dilutions plated non-selectively on YEPD and selectively on SC-Lys. Colonies were counted after two days and used for subsequent CO/NCO and sequencing analysis.

### CO-NCO phenotypic assay

Individual Lys^+^ colonies were inoculated into SC-Lys medium in 96-well plates and grown overnight (Hum and Jinks-Robertson 2019). The next day, 3 μl from each well were spotted onto YEPD plates and grown overnight. Cells were then replica plated onto SC-Lys, 5FOA (to distinguish COs from NCOs), and SC-Ura (to identify COs that are completely Ura^-^). Lys^+^ recombinants producing more than three papillae on 5FOA were scored as COs. To confirm CO assignment, 5FOA colonies from a given isolate were together inoculated into YEPD and grown overnight. Following DNA extraction, CO-specific primers (5’-CAGGTTTGTTCTGTCGAACG and 5’-TTGGTATGATTGCCCTTGGT) were used to amplify the 1.5 kb *hisG*-mediated deletion product on the putative V:II translocation product. Recombinants with confluent growth on 5FOA and no growth on SC-Ura were COs that had suffered an associated secondary HR event between the *hisG* repeats (Hum and Jinks-Robertson 2019).

### Sequencing and hetDNA analysis

Individual recombinants were inoculated into 96-well plates containing SC-Lys medium and genomic DNA was extracted after growth for at least 24 h. Each NCO was amplified using Phusion or ExTaq polymerase (New England BioLabs and Takara Bio, respectively) and a unique barcoded forward and reverse primer pair. Each primer contained complementarity to the recipient allele (forward 5’-ATGGTTGGGAAGTCATGGAAGTCG and reverse 5’-GCTTGGGAGTTGGGAATTGAAGTT) that was conjugated to a unique 16-nt barcode at its 5’ end (see https://www.pacb.com/products-and-services/analytical-software/multiplexing/ for Sequel system barcode sequences). Following amplification, PCR products were pooled in approximately equal concentrations into a single library (Gamble *et al*. 2019) and purified using the GeneJet PCR Purification Kit (Thermo Scientific). Ligation of SMRT bell adapters and amplicon sequencing using the PacBio Sequel system were done by the Duke Center for Genomic and Computational Biology. Following conversion of circular consensus sequence (CCS) reads to a FASTA format, sequences were analyzed using an in-house pipeline (Guo *et al*. 2015; Gamble *et al*. 2019). Instructions and program files for the pipeline are available at https://sites.duke.edu/jinksrobertsonlab/hetdna-mapping/. Only barcode pairs with at least 20 CCS reads and two distinct sequence species, with the minor one comprising at least 10% of the reads for the corresponding barcode pair, were included in data analyses. Recombinants that had only a single sequence species (i.e., no evidence of hetDNA) were excluded from analyses.

### Physical analysis of DSB induction and repair

DSB formation and repair were analyzed using Southern blots. Single colonies were inoculated into YEPR, diluted and grown overnight. At OD 0.7-1, I-*Sce*I or ZFN expression was induced with 2% galactose. Cells were isolated at the following times after galactose addition: 0 h, 0.75 h (45 min), 1.5 h, 4h, 6 h, and 12 h. Genomic DNA was then extracted and digested with *Hinc*II. Digests were run on 1% agarose gels at 4°. After electrophoresis, fragments were transferred from the gel to a charged nylon membrane (Roche). A *LYS2* probe was amplified and labeled with Digoxigenin (DIG)-dUTP using the primers 5’-TGAAGCCTTCCCAGAGAGAA and 5’-GCCAAGGAAAAATGTCTACCA (Roche PCR DIG Probe Synthesis Kit). The DIG probe was hybridized to the membrane (at 44°C; using Roche DIG Easy Hyb Granules for hybridization buffer) and detected using an anti-DIG antibody (Roche Anti-Digoxigenin-Ap Fav Fragments) and chemiluminescence film (Amersham Hyperfilm ECL). Following digestion, the uncut recipient and donor alleles produced ~2700-bp and ~3300-bp fragments, respectively. Following DSB formation the expected ~1400 bp fragment was very faint, which presumably reflected its rapid resection. Uncut fragment intensities were evaluated using ImageJ software (https://imagej.nih.gov/ij/) and the signal for the intact recipient allele was normalized to that of the donor allele at each time point. Each recipient:donor ratio was normalized to the ratio at the 0 h time point. The mean normalized ratios at each time point were plotted.

### Statistical Analysis

The proportions of CO-NCO products and the positions of hetDNA in NCO products were compared using Chi square contingency tests. For hetDNA length comparisons, the Mann-Whitney U test was used. For all tests, p<0.05 was considered significant. Repair frequencies and Southern blot recipient:donor ratios were deemed significant when 95% confidence intervals did not overlap.

### Data Availability

All strains are available upon request. Program files and a detailed protocol for the in-house pipeline used to analyze CCS reads from PacBio sequencing can be found on https://sites.duke.edu/jinksrobertsonlab/hetdna-mapping/. Raw CCS reads (fasta files) and barcodes of NCOs analyzed will be available on FigShare.

## RESULTS

Recombination studies to date have used either HO or I-*Sce*I, which each makes DSBs with 4-nt 3’ overhangs. We previously used I-*Sce*I to initiate HR between ectopic, 4.2 kb substrates (Guo *et al*. 2017; Hum and Jinks-Robertson 2019) and a modified version of this system was used in the current study (see below). To create DSBs with 4-nt 5’ overhangs on the ends, I-*Sce*I and its cleavage site were replaced by a zinc finger nuclease (ZFN) and its cleavage site. ZFNs designed to cleave the Drosophila *rosy* locus were used, with each component zinc-finger protein recognizing a 9-bp sequence flanking a 6-bp spacer (Beumer *et al*. 2006).

The effect of each end structure on HR intermediates and outcomes was examined using a common chromosomal assay based on *lys2* substrates (Figure 2A) (Guo *et al*. 2017; Hum and Jinks-Robertson 2019). To track the position and extent of donor sequence transferred during repair, single nucleotide polymorphisms (SNPs) were engineered into the donor allele at ~50 bp intervals (2% divergence). Mismatches within the resulting hetDNA were preserved by inactivating the *MLH1* gene, which encodes an essential component of the mismatch repair (MMR) machinery, and were detected by sequencing the *LYS2* allele in NCO products. The endogenous *LYS2* locus, located on chromosome II, contained the recipient allele where a galactose-inducible DSB was introduced into a region tolerant to amino acid changes. Insertion of the cleavage site for each enzyme created a −1 frameshift allele and resulted in a Lys^-^ phenotype. The donor repair template was a truncated *lys2* gene (*lys2Δ3’* inserted at the *CAN1* locus on chromosome V. To preserve homology with broken ends of the recipient allele, a corresponding cleavage-resistant site was derived by duplicating the 4-bp region normally flanked by enzyme-generated nicks (Figure 2B). Following DSB induction, copying of the additional 4 bp into the recipient corrects the −1 frameshift mutation and results in a selectable, Lys^+^ phenotype.

**Figure 2.**
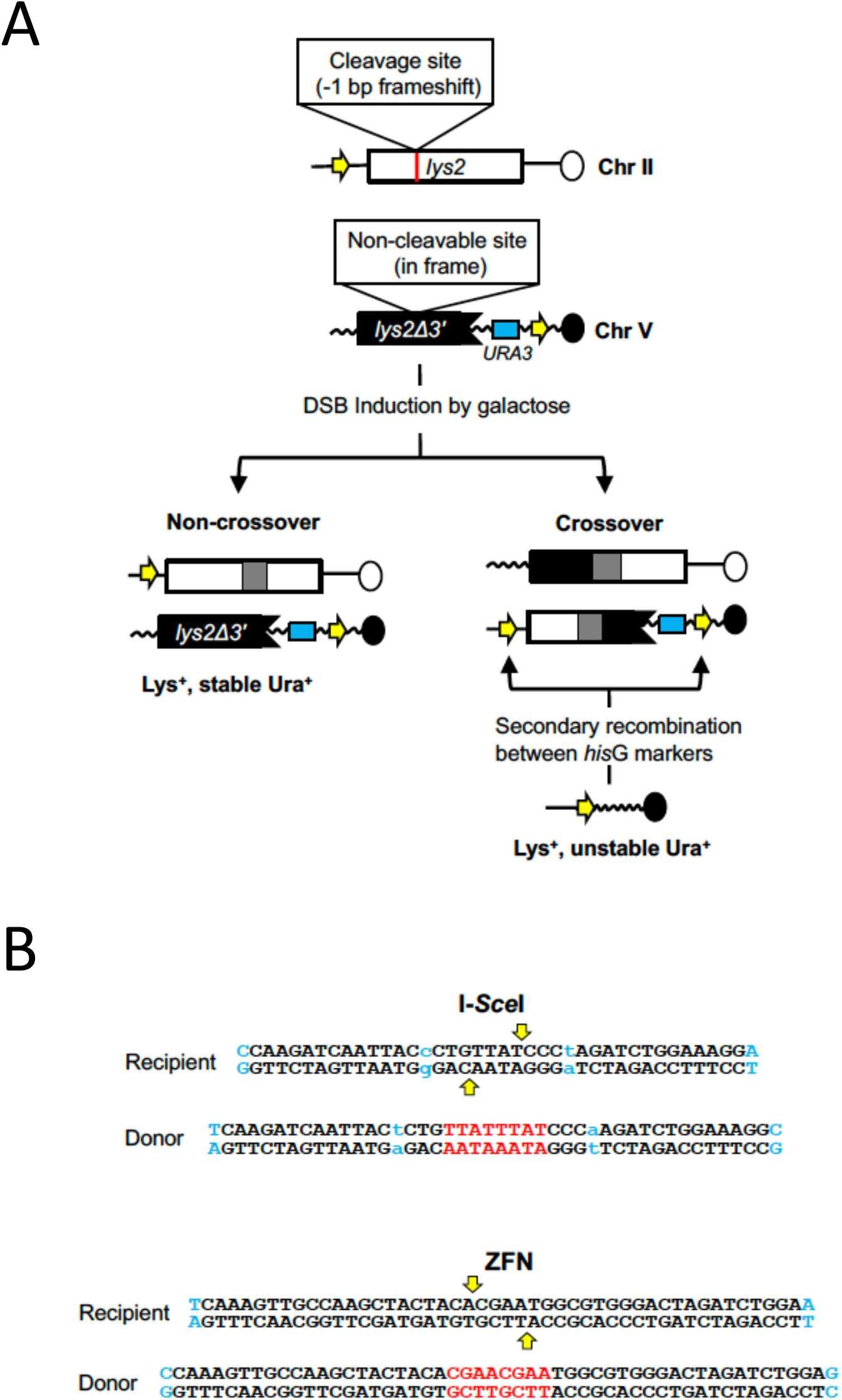
Ectopic recombination assay. (A) The recipient allele (white rectangle) is the endogenous *LYS2* gene on chromosome II. Insertion of a nuclease cleavage site into the recipient creates a −1 bp frameshift mutation. The donor (black rectangle) is a truncated *lys2* allele on chromosome V that contains a non-cleavable site created by duplicating 4 bp at the position of the recipient-allele break. Copying the additional 4 bp from the donor during repair corrects the frameshift mutation in the recipient allele. Gray rectangles in products represent hetDNA. The *hisG* (yellow arrows) and *URA3* (blue rectangle) markers were used to phenotypically distinguish between COs and NCOs. *hisG* direct repeats flank *URA3* in one of the CO products, resulting in an unstable Ura^+^ phenotype. (B) Sequences inserted into the recipient and donor alleles in each enzyme system are shown. Vertical yellow arrows represent the positions where the nucleases nick recipient DNA and the duplicated sequence in the donor is red. SNPs flanking the DSB are blue and those in the original I-*Sce*ISNP8 system are lowercase. While I-*Sce*I cleavage does not generate a gap relative to the donor allele, resection following ZFN cleavage creates an 8-bp gap. This small gap is not expected to affect repair efficiency.

The proportions of CO and NCO outcomes among Lys^+^ recombinants were determined using substrate-flanking markers that allow phenotypic discrimination (Hum and Jinks-Robertson 2019). In each enzyme system, a *hisG* marker was present upstream of the recipient allele and a *URA3-hisG* cassette was positioned downstream of the donor allele (Figure 2A). If an intermediate is repaired as an NCO product, then the *hisG* marker and *URA3-hisG* markers remain on separate chromosomes. If the intermediate is repaired as a CO product, however, then both *hisG* markers will reside on the same translocation product as direct repeats that flank the truncated *lys2* and *URA3* markers. A secondary recombination event between the *hisG* direct repeats results in frequent loss of the intervening segment containing the *URA3* marker, generating an unstable Ura^+^ phenotype detectable on 5FOA medium.

### DNA polymerase proofreading prior to 3’-end extension does not affect product profiles

In our previous studies using an I-*Sce*I-initiated DSB, the first SNP on each side of the break was positioned 8 nt from the resulting 3’ end in order to detect very short hetDNA tracts (I-*Sce*ISNP8) (Guo *et al*. 2017; Hum and Jinks-Robertson 2019). An unanticipated result was the frequent replacement of these SNPs with the donor SNPs, resulting in a small patch of apparent gene conversion on one or both sides of the DSB (Figure 3A). Among the 149 NCOs analyzed, 83% (124/149) lost the first SNP upstream of the DSB and 35% (52/149) lost the first SNP downstream of the break (Hum and Jinks-Robertson 2019). We further demonstrated that the proofreading activity of Pol δ was responsible for the efficient removal of the break-proximal SNPs (Guo *et al*. 2017). Because Pol δ backed up to remove the mismatch created by strand invasion/annealing before extending the end, it was possible that this influenced CO-NCO outcomes and/or affected hetDNA positions or lengths in NCO products.

**Figure 3.**
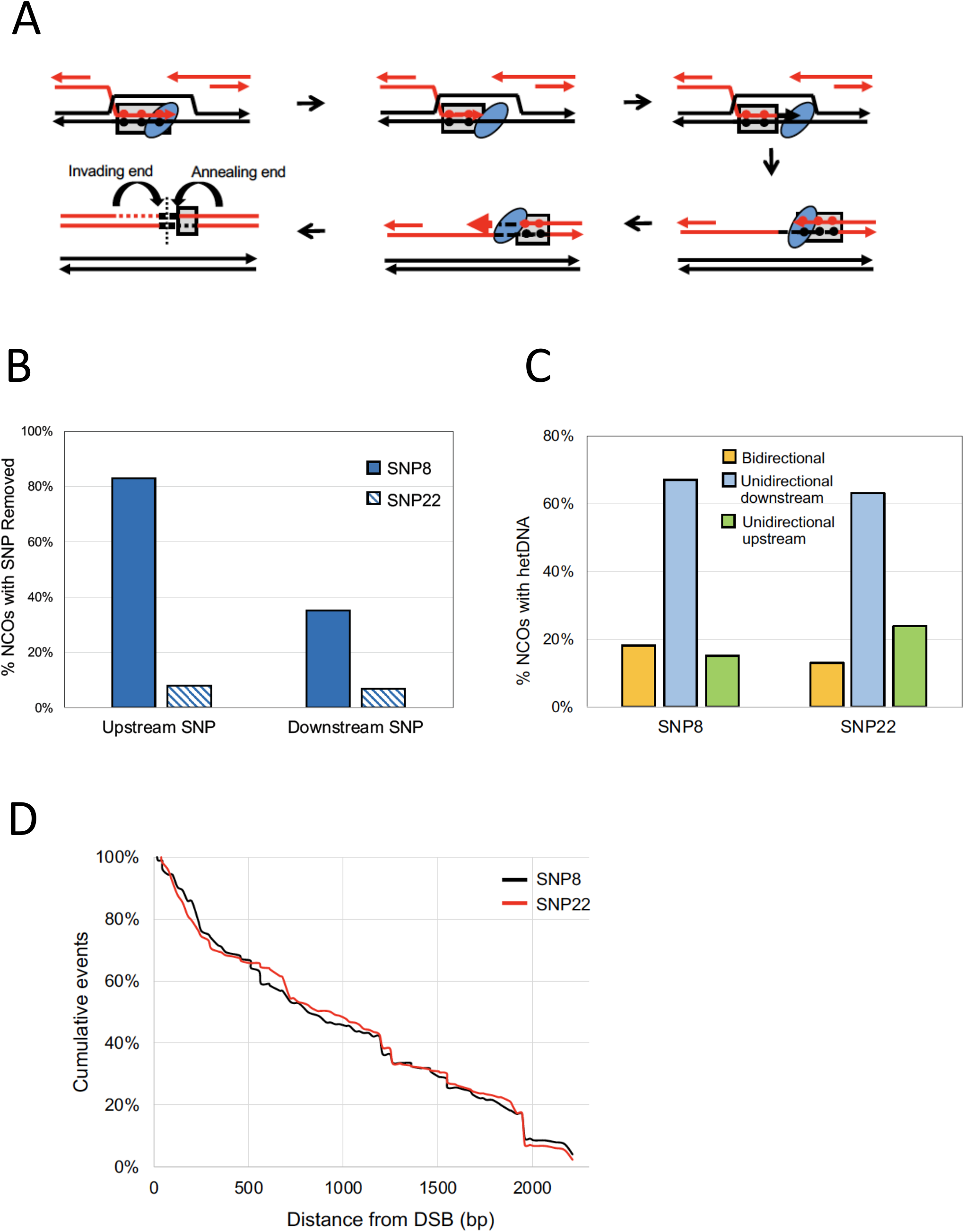
Effects of DNA polymerase proofreading activity on hetDNA profiles. (A) Upon strand invasion, hetDNA (gray boxes) is formed that contains mismatches between recipient and donor SNPs (red and black filled circles, respectively). DNA polymerase (blue oval) detects and excises the break-proximal mismatch *via* proofreading activity and then proceeds with DNA synthesis using the donor template. During SDSA, the newly synthesized 3’ end anneals to the other side of the break to create a new patch of hetDNA. Again, DNA polymerase recognizes and excises the terminal SNP created by annealing before continuing DNA synthesis. The resulting NCO product contains a small gene-conversion tract (black) on both sides of the break, followed by a single patch of hetDNA that marks the annealing end. The vertical dotted line indicates the position of the initiating DSB. Because proofreading reactions are independent, products may contain a short gene conversion patch on one side, both sides or neither side of the DSB. (B) Percentages of NCO hetDNA products that have the terminal upstream or downstream SNP removed. (C) Percentages of NCO products with bidirectional hetDNA tracts, unidirectional downstream tracts and unidirectional upstream tracts. (D) Cumulative distances of hetDNA lengths relative to the DSB at position 0.

The effect of terminal-SNP removal was examined by modifying the donor allele so that the first SNP on each side of the DSB was 22 nt instead of 8 nt from the 3’ end (I-*Sce*ISNP22). The increased distance of the SNPs from the ends should prevent their efficient proofreading by Pol δ (McCulloch *et al*. 2004; Anand *et al*. 2017; Guo *et al*. 2017) and as predicted, removal of the most break-proximal SNPs decreased when the distance from the 3’ end was increased (Figure 3B). With I-*Sce*ISNP22, removal of the terminal SNP upstream of the DSB occurred only 8% of the time (23/277; p<0.001); downstream of the DSB only 7% (19/277; p<0.001) of NCOs had lost the recipient SNP.

In previous analyses with the I-*Sce*ISNP8 system, 7% (33/471) of Lys^+^ products were CO events (Hum and Jinks-Robertson 2019). This distribution did not change significantly with the I-*Sce*ISNP22 substrates, where 9% (35/372) of products were COs (p=0.25). Among NCOs, hetDNA tracts were classified based on their position relative to the initiating DSB (Figure 3C). hetDNA tracts on only one side of the break are characteristic of the SDSA pathway (Figure 1) and were either unidirectional upstream (promoter proximal) or unidirectional downstream (promoter distal). The third, bidirectional category of NCOs contained hetDNA on each side of the DSB and is consistent with either Holliday junction dissolution or a double SDSA event. With the I-*Sce*ISNP8 substrates, 18% (27/149) of hetDNA tracts were bidirectional and among the unidirectional tracts, 82% (100/122) were downstream of the initiating DSB (Hum and Jinks-Robertson 2019). Moving the first SNP to 22 nt from the 3’ end did not alter these distributions; 13% (35/277) of hetDNA tracts were bidirectional (p=0.1) and of the unidirectional tracts, 73% (176/242) were downstream of the DSB (p=0.07) (Figure 3C; complete hetDNA profiles are in Figure S1). Lastly, the distributions of hetDNA lengths for the I-*Sce*ISNP8 *versus* I-*Sce*ISNP22 substrates were compared (Figure 3D). For this analysis individual tracts on each side of the DSB were compiled into single data sets and plotted as cumulative distances from the break. The median hetDNA length was 896 bp and 922 bp, respectively for the I-*Sce*ISNP8 and I-*Sce*ISNP22 substrates (p=0.95, Mann-Whitney U Test). These results indicate that engaging the proofreading activity of Pol δ prior to end extension has no discernable effect on HR intermediates or outcome. In experiments using the ZFN to initiate HR, the first SNP was 22 nt from each of the 3’ ends created by enzyme cleavage (Figure 2B).

### Effect of initiating enzyme on HR outcomes

We analyzed at least 300 Lys^+^ recombinants resulting from each type of initiating DSB (Figure 4A). Among products of I-*Sce*I-initiated events (3’ overhangs), 9% (35/372) were COs and 91% (337/372) were NCOs. Following initiation with ZFN there was a modest increase in the proportion of COs to 15% (55/377) (p=0.04 by Chi square test).

**Figure 4.**
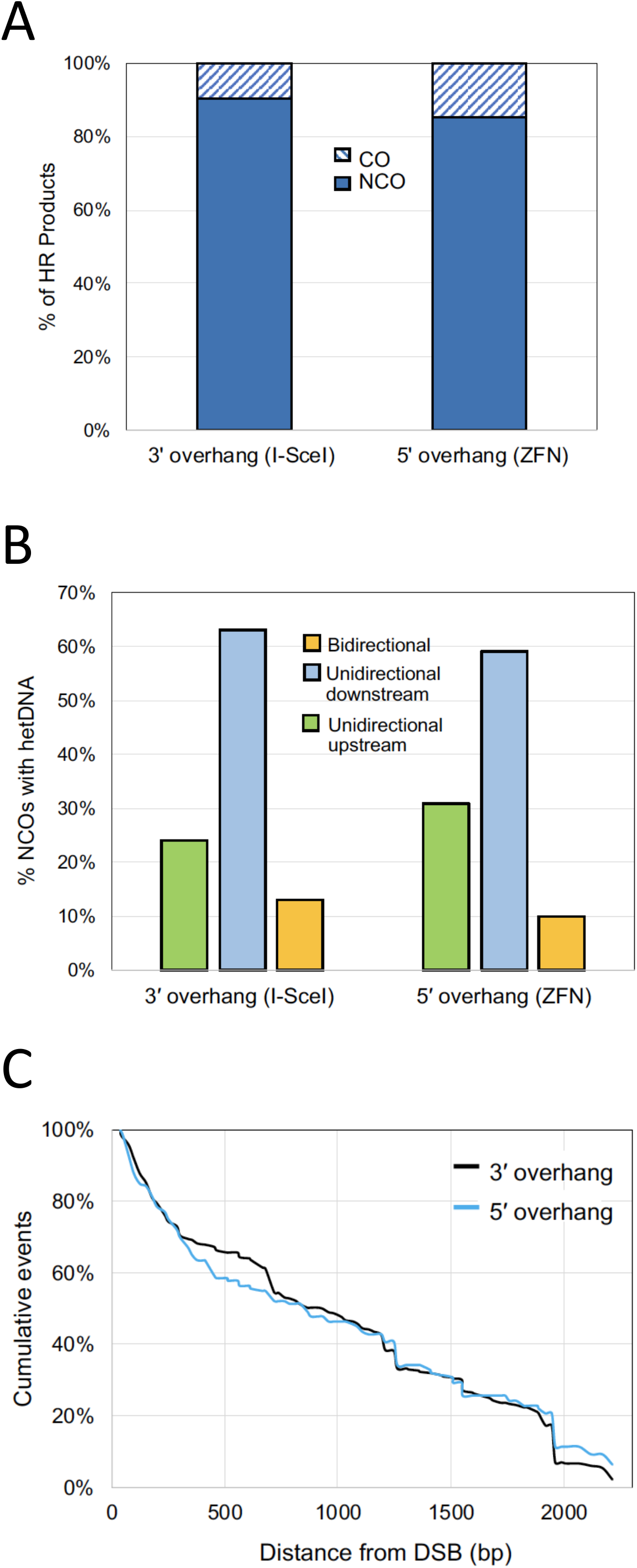
Effect of initiating enzyme on repair outcomes. (A) Percentage of HR products that are COs or NCOs for each type of DSB. (B). Percentage of NCO products with bidirectional hetDNA tracts, unidirectional downstream tracts and unidirectional upstream tracts for each type of DSB. (C) Cumulative hetDNA tract lengths relative to the initiating DSB at position 0.

The sequencing profiles of NCOs derived from I-*Sce*I or ZFN cleavage are presented in Figure S1. Among the NCO products derived from an I-*Sce*I generated DSB, 13% (35/277) had bidirectional hetDNA; of the unidirectional tracts, 73% (176/242) were downstream and 27% (66/242) were upstream of the break, respectively (Figures 4B). For NCOs initiated by the ZFN, 10% (13/127) had bidirectional hetDNA; 66% (75/114) and 34% (39/114) of unidirectional tracts were downstream and upstream of the DSB, respectively. There was no difference in unidirectional versus bidirectional hetDNA (p=0.60) or in the position of unidirectional tracts relative to the initiating break (p=0.22). The hetDNA tract lengths relative to each type of initiating DSB are plotted in Figure 4C. I-*Sce*I-generated DSBs with 3’ overhangs and ZFN-generated DSBs with 5’ overhangs had a similar median hetDNA lengths: 922 bp (black line) and 832 bp (blue line), respectively (p=0.1).

### Efficiency of nuclease-generated DSBs

In the *mlhlΔ* background the repair (Lys^+^) frequencies for DSBs with 3’ overhangs (I-*Sce*I) and 5’ overhangs (ZFN) following the addition of 0.1% galactose for 45 min were 0.0471 and 0.191, respectively. This difference could be due to more efficient repair of ZFN breaks or more efficient cleavage by ZFN. Southern blot analysis was used to physically monitor the formation and repair of DSBs with 3’ overhangs *vs* 5’ overhangs (Figure 5A-B). Because this analysis required maximal enzyme induction (2% galactose), we also measured the repair frequency when cells were plated on 2% galactose medium. After constitutive DSB induction the repair frequencies were 0.46 for I-*Sce*I-induced breaks and 0.82 for DSBs with a 5’ overhang created by ZFN. In Southern analyses, the loss and reappearance of the full-length recipient fragment was monitored after DSB induction by either I-*Sce*I or ZFN, and was normalized to the donor-allele fragment, which remained constant throughout the experiment (Figure 5C). Samples were taken at 0 h, 0.75 h, 1 h, 1.5 h, 3 h, 6 h, and 12 h after galactose addition. The maximum level of the cut recipient fragment occurred at 3 h with both I-*Sce*I and ZFN, but more efficient cleavage occurred with ZFN. ZFN cleaved 82% of recipient alleles while I-*Sce*I cleaved only 48%; repair for both was complete by 24 h. The observed cleavage difference is consistent with the 2-fold difference in repair efficiencies. Interestingly, between 3h and 6h after galactose addition, 16% of ZFN-induced breaks were repaired while 10% of I-SceI-induced DSBs were repaired. These data indicate that ZFN more efficiently generated DSBs, and that DSBs with 5’ overhangs are repaired somewhat faster than I-SceI-induced breaks.

**Figure 5.**
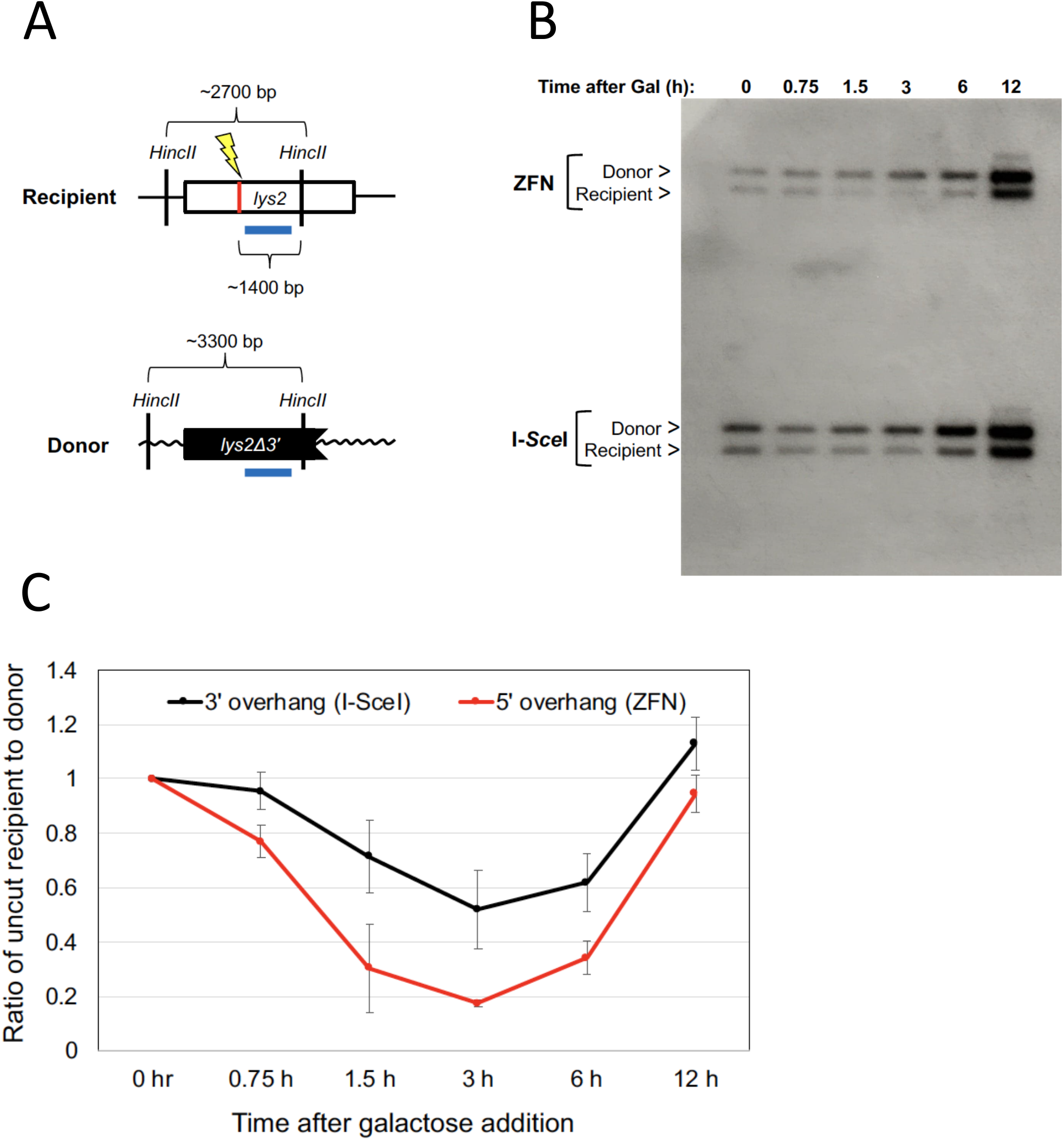
Physical analysis of DSB formation and repair. (A) Schematic of fragment sizes expected following cleavage of genomic DNA with *Hinc*II; the blue bar represents the position of the probe hybridized to Southern blots. (B) Samples were taken at the indicated times after enzyme induction with 2% galactose. Positions of donor and intact recipient fragments are indicated. (C) The average ratio of the intact recipient fragment remaining at each time point relative to the donor fragment was calculated from four independent inductions. The recipient/donor ratio in each was normalized to the ratio at t=0. The 0.75 h and 12 h mean values were analyzed in only two of the four experiments.

## DISCUSSION

Mitotic HR mechanisms/pathways have largely been defined by following repair of a sitespecific DSB. Studies to date, including our own, have used either I-*Sce*I or HO, each of which generates 4-nt 3’ overhangs (Bellaiche *et al*. 1999; Guo *et al*. 2017; Haber 2000; Hum and Jinks-Robertson, 2017, 2018, 2019; Katz *et al*. 2014; Pâques and Haber, 1999; Sugawara and Haber 2006). Here, we used a ZFN that creates 4-nt 5’ overhangs to examine whether the source of the initiating break is of any functional consequence in terms of HR outcomes and molecular intermediates. We additionally examined whether degradation of the invading/annealing 3’ end by Pol δ-mediated proofreading (Guo *et al*. 2017) affects subsequent DNA synthesis.

In the first iteration of our I-*Sce*I system, the DSB-proximal SNPs were only 8 nt from the enzyme-generated 3’ ends (I-*Sce*ISNP8), which resulted in their frequent removal by proofreading (Guo *et al*. 2017). Because the D-loop formed upon strand invasion is a highly dynamic structure (Piazza *et al*. 2019), a proofreading-related delay in the initiation of end extension might limit the extent of DNA synthesis prior to D-loop collapse and/or affect second-end capture. Such a scenario could, in principle, explain why hetDNA tracts in our ectopic assay are much shorter than those associated with spontaneous or DSB-induced HR in diploids (Hum and Jinks-Robertson 2019; Yin *et al*. 2017). Although moving the DSB-proximal SNPs from 8 nt to 22 nt from the 3’ ends significantly decreased their loss, there was no difference in the CO-NCO distribution of HR products or the positions/lengths of NCO-associated hetDNA. Similar, Pol δ-mediated 3’-end degradation has been observed in a break-induced replication (BIR) assay (Anand *et al*. 2017), suggesting that exonucleolytic removal of break-proximal mismatches is a general feature of yeast recombination. In contrast to the BIR assay, however, we did not detect processive degradation of ends by Pol δ in our prior studies (Guo *et al*. 2017). This could reflect the greater distance between SNPs in our system, which are ~50 bp apart, or the very short regions of homology (108 bp) used in the BIR system. Although we assume that the 3’-end degradation was mismatch-triggered, an interesting possibility is that 3’-end excision by Pol δ is a general feature of DNA synthesis that initiates in the context of recombination.

DSBs created by a ZFN were associated with proportionally more COs than were I-*Sce*I-generated DSBs, but there was no difference in either the position or length of hetDNA in NCO products. Whether the CO-NCO difference reflects the nature of the enzyme-generated ends or some other property of the enzymes (e.g., different dissociation kinetics or asymmetry in enzyme binding) cannot be distinguished. In an earlier study of MMR-mediated antirecombination during repair of I-*Sce*I-induced DSBs, we found that changes in HR product distribution correlated with changes in the positions of hetDNA in NCO products (Hum and Jinks-Robertson 2019). Specifically, an increase in CO frequency was accompanied by an increase in the frequency of bidirectional hetDNA in NCO products, each of which requires engagement of the donor by both broken ends. Although these data appear to be contradictory to that reported here, it should be noted that the previous correlation between COs and bidirectional hetDNA reflected features of MMR-mediated anti-recombination and was seen specifically in relation to disabling mismatch-binding versus mismatch-processing activity.

Induction of the ZFN was associated with 2-fold more recombinants than was I-*Sce*I induction and this correlated with the relative cleavage efficiencies of the enzymes. In addition to the differing cleavage efficiencies, there appeared to be a difference in DSB repair rates, with ZFN breaks being repaired faster than those created by I-*Sce*I. This difference could reflect relatively slow release of I-*Sce*I from ends, which would delay the initiation of resection, or specific end preferences of DSB-binding proteins. In NHEJ studies, for example, Ku was recruited much more rapidly recruited to ZFN-generated ends than to ends created by HO (Liang *et al*. 2016). An interesting possibility is that rapid end binding might result in more coordinated resection of ends, which in turn might alter HR outcomes, as observed here. *In vitro* studies using *E.coli* or yeast proteins have shown that the recruitment and activity of resection enzymes also differ based on DSB-end structure (Morimatsu and Kowalczykowski 2014; Cannavo *et al*. 2013), and this could potentially affect molecular intermediates. Although we found no significant difference in the lengths of hetDNA tracts associated with an I-*Sce*I-*versus* a ZFN-initiated DSB, a wild-type background in which all resection activities were present was used. End structure preferences might affect only the early stages of HR, with steps following the initiation of 5’ end resection being unaffected and leading to similar repair outcomes, regardless of the initiating break. Removal of Ku, MRX, Exo1 or Sgs1 may reveal differences due to end-structure preferences of the corresponding complexes.

## ACKNOWLEDGEMENTS

We would like to thank Dana Carroll for the generous contribution of ZFN plasmids. We also acknowledge Yee Fang Hum for construction of plasmids and strains used in this work. Finally, we want to thank Tom Petes and other members of the Jinks-Robertson lab for valuable feedback during the course of this project and comments on the manuscript.

---

**Figure S1.**
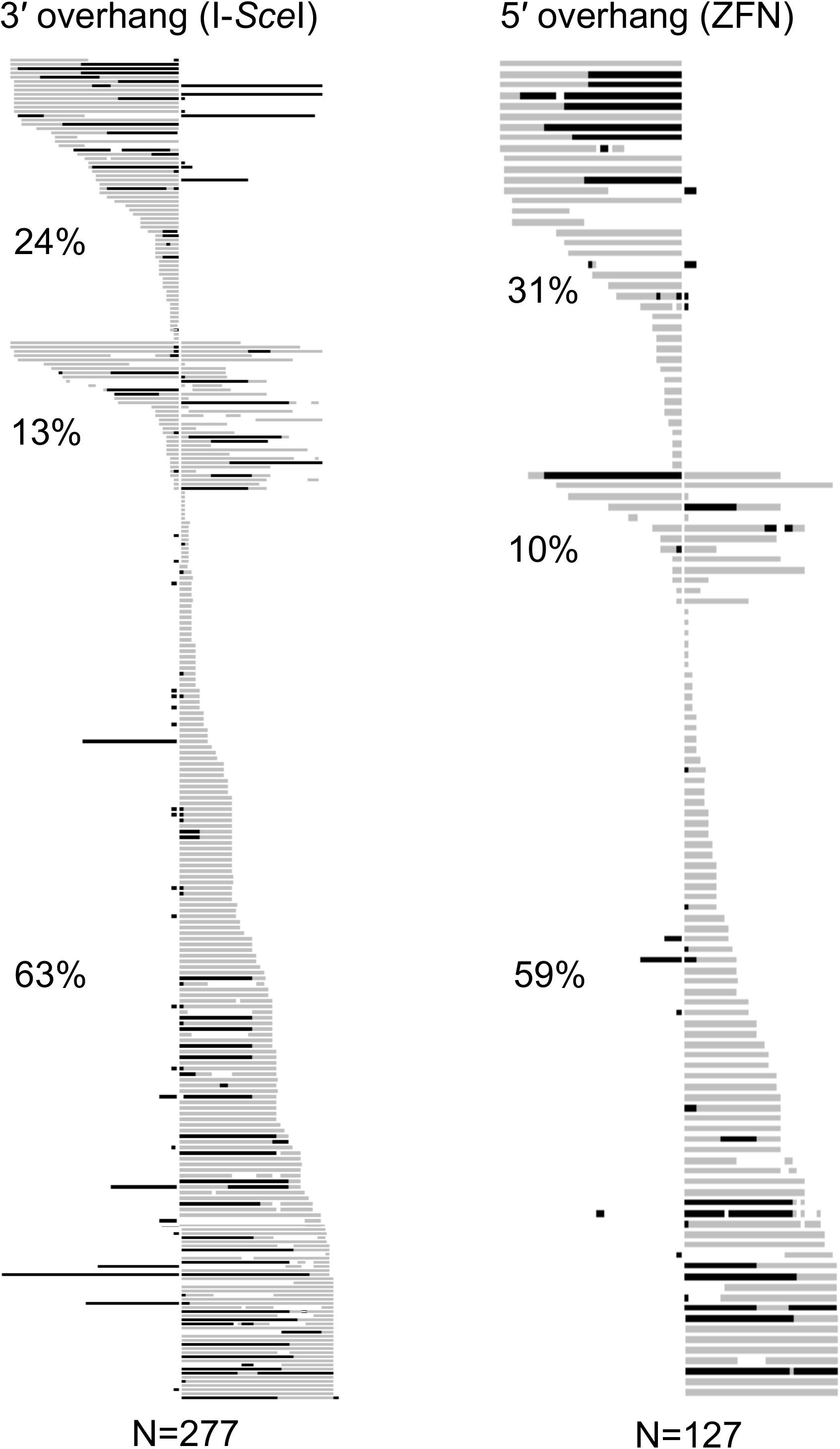
hetDNA profiles in NCO products. Each horizontal line represents an individual NCO product and the vertical white line indicates the DSB position. Products are grouped by hetDNA position. Black areas signify the presence of only donor SNPs, which reflect regions of gene conversion or gap expansion, while gray areas represent hetDNA tracts containing both donor and recipient SNPs. White regions flanked by black and/or gray indicate the presence of only recipient SNPs and could reflect restoration that is independent of MMR, simultaneous invasion of more than one repair template, or template switching (see Hum and Jinks-Robertson 2019 for a discussion).

**Table S1.**
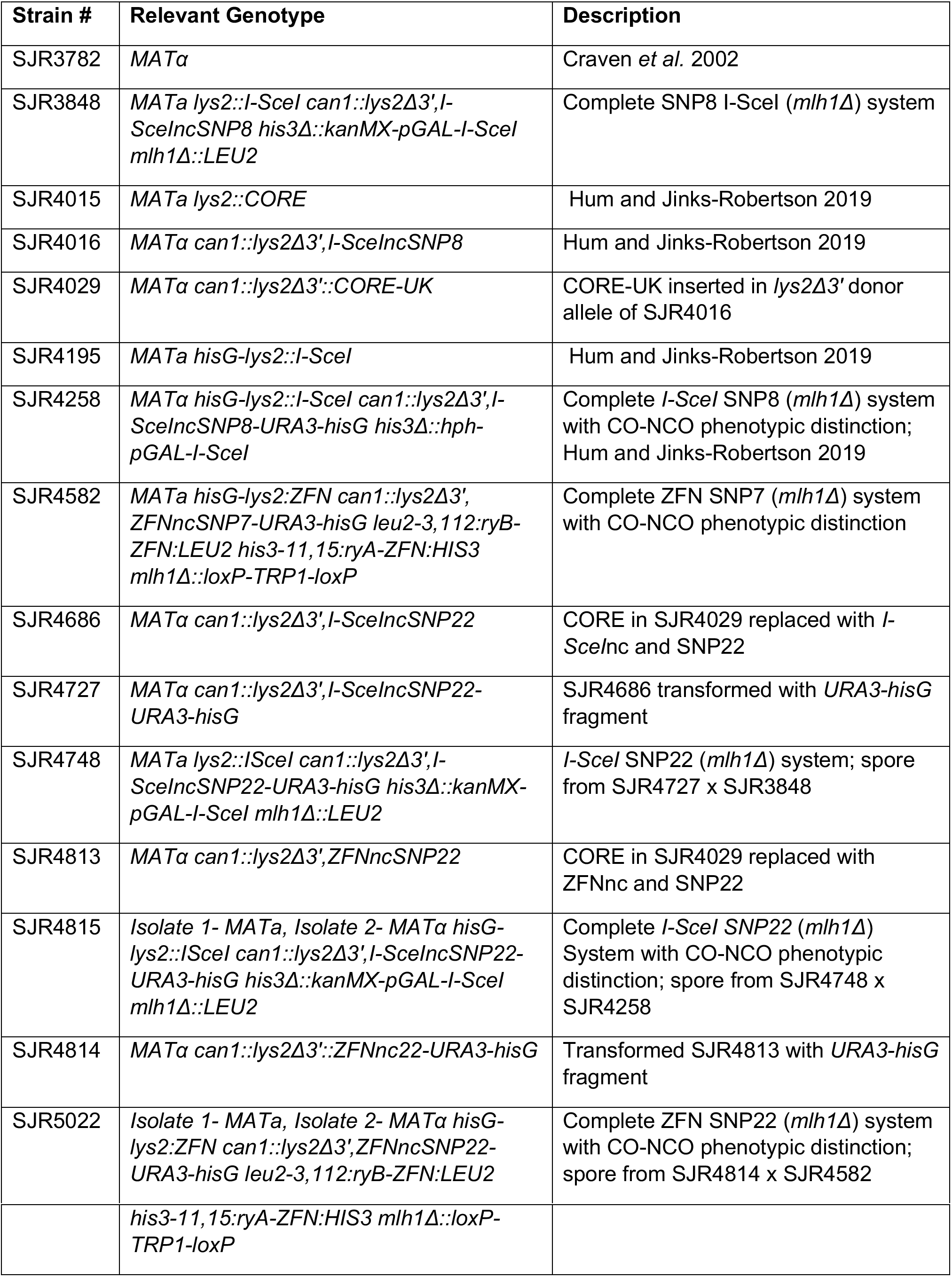
Yeast Strains

**Table S2.** Plasmids

| **Plasmid #** | **Relevant Info** | **Reference** |
| --- | --- | --- |
| pNY51 | *hisG-URA3-hisG* cassette | Alani *et al.* 1987 |
| pRS303 | Ylp vector containing *HIS3* | Sikorski and Hieter 1989 |
| pRS305 | Ylp vector containing *LEU2* | Sikorski and Hieter 1989 |
| *ryB*-ZFN | ZFN construct | Beumer *et al.* 2006 |
| *ryA*-ZFN | ZFN construct | Beumer *et al.* 2006 |
| pSR1109 | *GAL1* regulated ZFN *ryA* fragment cloned into digested pRS303 |  |
| pSR1110 | *GAL1* regulated ZFN *ryB* fragment cloned into pRS305 |  |

